# Evidence for a hyper-reductive redox in a sub-set of heart failure patients

**DOI:** 10.1101/246413

**Authors:** Sairam Thiagarajan, Amit N. Patel, Meenu Subrahmanian, Rajendran Gopalan, Steven M. Pogwizd, Sudha Ramalingam, Sankaran Ramalingam, Namakkal S. Rajasekaran

**Affiliations:** Cardiac Aging & Redox Signaling Laboratory, Molecular and Cellular Pathology, Department of Pathology/Center for Free Radical Biology, University of Alabama at Birmingham, Birmingham, AL, USA; PSG Center for Molecular Medicine and Therapeutics, PSG Institute of Medical Sciences & Research (Affiliated to the Tamilnadu Dr MGR Medical University), Coimbatore, Tamil Nadu, India; Division of Cardiothoracic Surgery, University of Miami – Miller School of Medicine, FL, USA; Department of Cardiology, PSG Institute of Medical Sciences & Research (Affiliated to the Tamilnadu Dr MGR Medical University), Coimbatore, Tamil Nadu, India; Comprehensive Cardiovascular Center, Department of Medicine, University of Alabama at Birmingham, Birmingham, AL, USA; Cardiovascular Medicine, University of Utah School of Medicine, Salt Lake City, UT, USA

**Keywords:** hyper-reductive, hyper-oxidative, heart failure, circulatory redox state

## Abstract

**Background:** Oxidative stress has been linked to heart failure (HF) in humans. Antioxidant-based treatments are often ineffective. Therefore, we hypothesize that some of the HF patients might have a reductive stress (RS) condition. Investigating RS-related mechanisms will aid in personalized optimization of redox homeostasis for better outcomes among HF patients.

**Methods:** Blood samples were collected from HF patients (n=54) and healthy controls (n=42) and serum was immediately preserved in –80°C for redox analysis. Malondialdehyde (MDA; lipid peroxidation) levels by HPLC, reduced glutathione (GSH) and its redox ratio (GSH/GSSG) using enzymatic-recycling assay in the serum of HF patients were measured. Further, the kinetics of key enzymatic-antioxidant enzymes was analyzed by UV-Vis spectrophotometry. Non-invasive echocardiography was used to relate circulating redox status with cardiac function and remodeling.

**Results:** The circulatory redox state (GSH/MDA ratio) was used to stratify the HF patients into normal redox (NR), hyper-oxidative (HO), and hyper-reductive (HR) groups. While the majority of the HF patients exhibited the HO (42%), 41% of them had a normal redox (NR) state. Surprisingly, a subset of HF patients (17%) belonged to the hyper-reductive group, suggesting a strong implication for RS in the progression of HF. In HF patients, SOD, GPx and catalase were significantly increased while GR activity was significantly reduced relative to healthy controls. Furthermore, echocardiography analyses revealed that 55% of HO patients had higher systolic dysfunction while 75% of the hyper-reductive patients had higher diastolic dysfunction.

**Conclusion:** These results suggest that RS may be associated with HF pathogenesis for a subset of cardiac patients. Thus, stratification of HF patients based on their circulating redox status may serve as a useful prognostic tool to guide clinicians designing personalized antioxidant therapies.

## Background

There is a general consensus that oxidative stress induces various pathophysiological processes including cardiovascular complications [1, 2]; however, counteracting antioxidant supplementations have failed to prevent the progression or curtail disease pathogenesis [3, 4]. At present, it is not clear whether oxidative stress is a cause or consequence in a given cell or organ system exhibiting a chronic disease state. Therefore, it is vital to critically analyze the global redox milieu of patients experiencing chronic illnesses including heart failure (HF). HF is a progressive condition in which cardiac muscle weakens and becomes inefficient to meet the body’s demand for blood and oxygen supply. The etiology of HF is multifaceted as several genetic, biochemical, electrical and inflammatory factors have been shown to underlie the structural and functional remodeling that develops over time [5-8]. Based on the currently available literature, a majority of the HF conditions have been correlated with oxidative stress for the past several decades. In particular, ischemic heart disease and/or reperfusion injury have been shown to display a hyper-oxidative state wherein increased reactive oxygen and nitrogen species (ROS/RNS) generation correlates with a worsening of myocardial injury [9-11]. In spite of these observations, supplementation with antioxidants seems to be inefficient to treat such conditions in a failing heart [12-14]. In particular, pre-clinical observations using rodent models have documented that a forced induction of oxidative stress leads to “heart failure” and pre-treatment with potential antioxidants seems to be protective [15-17]. However, these findings were not reproducible in HF patients [18-20].

To the best of our knowledge, all HF studies have focused on documenting the differences between HF patients and healthy control groups, and have not examined the potential for individual variations in the context of redox status among HF patients. Importantly, findings based on a group may not be precise to each individual of that group. Therefore, considering the inconsistent effects of antioxidant trials in human patients, it is worth testing whether all HF patients experience similar redox state. The ROS/RNS produced during basal mitochondrial metabolism (oxygen consumption at resting state) or in response to physical activity are key modulators of cellular motility to maintain a redox homeostasis and preserve the dynamic function of the myocardium [21-24]. However, other factors including genetic or chronic stresses that modulate ROS/RNS may tip the redox milieu towards either direction of the redox spectrum (i.e. reductive or oxidative). Despite several studies demonstrating the futility and/or detrimental effects of antioxidants, there has not been a single study attempting to understand the mechanisms associated with failure of the antioxidants in over six decades of biomedical research. In the present study, we attempt to address this critical gap in knowledge and postulate that some HF patients may either exhibit a hyper-reductive or normal redox state potentially conferring vulnerability and detrimental side effects to antioxidant treatment.

In the current study, we determined the circulatory redox state of HF patients by measuring glutathione redox (GSH:GSSG) ratio, lipid peroxidation (MDA) and the activities of key antioxidant enzymes to compare with the healthy control group. Moreover, we utilized the ratio of GSH/MDA as a peripheral redox index and focused on stratifying HF patients according to this measure. Our findings revealed a surprising but meaningful observation in that there appears to be distinct subsets of HF patients exhibiting divergent redox signatures. Therefore, our data indicate that not all HF patients have oxidative stress as traditionally reported. Our pilot observations in this small cohort of HF patients (n=54) warrant a new redox-based classification in these patients for selecting an appropriate therapy.

## Methods

### Study population

Our study population included heart failure (HF) patients (n=54) and healthy controls (n=42) who attended the in-patient and out-patient department of cardiology in PSG Hospitals, Tamil Nadu, India. Their characteristics are tabulated in Table 1 and 2. Patients with HF who were diagnosed using WHO criteria were considered for this study. Patients on dialysis, as well as those exhibiting severe liver diseases, malignancies or consuming antioxidant supplements were excluded from the study. While 85% of patients were admitted to the hospital for the first time, the remaining were presented for follow-up visits. Conventional HF therapy was given to all patients including angiotensin-converting enzyme inhibitors (ACE) inhibitors, beta blockers, and angiotensin receptor blockers according to guidelines. The study was approved by the PSG Institutional Human Ethics Committee (IHEC) and all patients/subjects completed a written informed consent prior to their participation.

**Table 1:** Demographic details of Cardiac failure patients.

| Characteristics of patients | Cardiac failure patients ( n=54) |  |  |
| --- | --- | --- | --- |
|  | Hyperreductive<br>(n=9) | Hyperoxidative<br>(n=23) | Normal redox<br>(n=22) |
| **Age in years** : Mean age(Range) | 50 (32 - 75) | 58 (34 - 73) | 52 (34 - 75) |
| **Sex** : n (% male) | 9 (100) | 23 (100) | 21 (95) |
| **Mean Height in cm (Range)** | 164 (151.7 -177 ) | 163 (152 - 173) | 163 (143 - 178) |
| **Mean Weight in kg (Range)** | 60 (50 - 70) | 62 (41 - 75) | 63 (38 - 100) |
| **Body mass index (kg/m2) (Range)** | 22 (19.5 - 24.5) | 23 (16.4 - 30.3) | 24 (15.2 - 35.4) |
| **Hemodynamics** |  |  |  |
| Heart rate (No. of times/ minute) | 97 (60 - 100) | 94 (74 - 121) | 96 (70 - 116) |
| Mean Systolic Blood pressure (mmHg) | 129 (80 - 170) | 129 (80 - 180) | 125 (90 - 170) |
| Mean Diastolic Blood pressure (mmHg) | 79 (60 - 140) | 85 (60 - 110) | 77 (50 - 110) |
| **Comorbidities ,%** |  |  |  |
| DM & DM assoiated disorders | 55 | 65 | 45 |
| Systemic Hypertension | 33 | 22 | 27 |
| Hypothyroidism | - | 17 | 4.5 |
| Chronic kidney Disease | 11 | 13 | - |
| Anaemia | - | 7 | 9 |
| BPH (Benign Prostatic Hyperplasia) | 11 | - | 4.5 |
| Hepatitis | - | - | 4.5 |
| CVA (Cerebrovascular Accident) | - | 4 | - |
| COPD | 11 | 9 | 14 |
| **Cardiac Disease Pathology, %** |  |  |  |
| Dilated Cardiomyopathy | 55 | 35 | 41 |
| Heart Failure | 78 | 52 | 77 |
| Coronary Artery Disease | 22 | 43 | 45 |
| LV Dysfunction | 78 | 52 | 68 |
| Rheumatoid Heart Disease | 11 | - | 9 |
| Ischemic Heart Disease | - | 4 | 4.5 |
| Aorto Iliac disease | 11 | 22 | - |
| Biventricular dysfunction | - | 4 | 4.5 |
| **Medication, %** |  |  |  |
| Diuretics | 55 | 87 | 82 |
| β blockers | 44 | 52 | 54 |
| Statins | 11 | 39 | 27 |
| ACE Inhibitors | 33 | 43 | 45 |
| Digitalis glycosides | 33 | 48 | 18 |
| Anticoagulant | 11 | 22 | 9 |
| (ARB) angiotensin II antagonists. | - | 22 | 4.5 |
| Antiplatelet medications | - | 22 | 14 |

**Table 2:** Demographic details of healthy controls

| **Characteristics of patients** | **Healthy controls (n=42)** |
| --- | --- |
| **Age in years** : Mean age(Range) | 36.35 ± 12.2 (18-64) |
| **Sex** : n (% male) | 27 (64) |
| **Mean Height in cm** (Range) | 165.8 ± 9 (148 - 192) |
| **Mean Weight in kg** (Range) | 62.7 ± 11 (38 - 89) |
| **Body mass index** (kg/m<sup>2</sup>) (Range) | 22.9 ± 4.2 (16.1 - 34.6) |
| **Hemodynamics** |  |
| Heart rate (No. of times/ minute) | 78 ± 9 (64 - 98) |
| Mean Systolic Blood pressure (mmHg) | 119 ± 17 (90 - 190) |
| Mean Diastolic Blood pressure (mmHg) | 76 ± 11 (50 - 110) |

### Reagents

Supplies and reagents were purchased from Sisco Research Laboratories (India) unless otherwise specified.

### Clinical assessment and echocardiography

Validation and confirmation of HF in patients was made based on clinical and echocardiography assessment. One echocardiogram was selected for each patient at the time of enrollment. Two-dimensional and M-mode echocardiography, color flow and spectral Doppler as well as annular TDI data were obtained from all patients and healthy controls using an ultrasound system (Philips IE33, Netherlands). Standard views, including the parasternal long axis view, short axis at the papillary muscle level and apical four and two chamber views were recorded. Cardiac chamber dimensions, volumes and left ventricular mass were measured according to current recommendations [25]. Left ventricular ejection fraction (LV EF) was calculated using the teichholz formula. Mitral Doppler signals were recorded in apical 4 chamber view, with Doppler sample volume placed at the tip of the mitral valve leaflets. The following parameters were obtained: early diastolic mitral inflow peak velocity (E), late diastolic mitral inflow peak velocity (A) and their ratio.

### Blood sampling

Blood samples were obtained by venipuncture using a 21-gauge needle from the patients and healthy control subjects in plain tubes. Serum was separated by centrifugation at 3000 rpm for 5 minutes at 4ºC and immediately aliquoted and frozen at −80ºC. The stored aliquots of serum were used for analyzing redox status and the activities of antioxidant enzymes.

### Lipid peroxidation

Serum malondialdehyde levels were measured by the DNPH (2,4 dinitrophenylhydrazine) derivatization method [26]. Briefly, 125μl of serum was diluted in 125 μl 1XPBS (pH 7.4), mixed with 50μl of 6M NaOH and incubated at 60°C for 30 minutes. 125μl of 35% perchloric acid was then added to the mixture. 250μl of this mixture was combined with 25μl of 5mM DNPH (in 2M HCl) and incubated in the dark for 30 minutes. An aliquot of 25μl of the solution was injected into the HPLC system (Shimadzu, Japan). The mobile phase of 0.2% (v/v) acetic acid and acetonitrile (50: 50 v/v) was run at a flow rate of 0.5ml/ minute at 25°C. A C18 column (Agilent, USA) was used and the chromatograms were obtained at 310 nm. Concentration of MDA was measured after comparing with a reference curve using TMP (1, 1, 3, 3 - Tetramethoxypropane) as a standard.

### Reduced glutathione levels

The spectrophotometric based glutathione reductase/5, 5’-dithio-bis (2-nitrobenzoic acid) (DTNB) recycling assay was used to measure reduced glutathione (GSH) levels as per [27-29] with minor modifications. In short, metaphosphoric acid extracts of serum samples were prepared and treated with triethanolamine (TEAM reagent) to adjust the pH for total GSH quantification according to the manufacturers’ protocols (GSH redox Kit #CS0260, Sigma-Aldrich). An aliquot of TEAM treated samples were mixed with 1.0 mM 2-vinyl pyridine and incubated for 1.0 hour at room temperature for GSSG measurements. For enzymatic recycling, the processed samples were treated with a reaction mixture containing 0.25mM dithiobis-nitrobenzoic acid (DTNB), 0.38U/ml Glutathione reductase and 60μl of 0.17mM NADPH in 0.1M sodium phosphate buffer (pH 7.4) with 5mM EDTA. The rate of DTNB formation was measured at A412 nm. Similarly, GSH and GSSG standards were treated and measured to obtain a standard graph to extrapolate the values from serum samples. The concentration of reduced glutathione (GSH) was estimated by subtracting the measured oxidized (GSSG) glutathione levels from the measured total (GSH plus GSSG) glutathione. GSH/GSSG ratio was then determined.

### SOD activity

Superoxide dismutase activity in the serum samples were measured using the pyrogallal auto-oxidation method [30]. This assay was carried out in a microtiter plate and the readings were measured using multimode reader (Varioskan Flash, Thermoscientific, USA). Approximately 50μl of serum was treated with 25μl ethanol and 15μl chloroform. The contents were vortexed and centrifuged at 8000 rpm for 5 minutes and the supernatant was analyzed for SOD activity by mixing with 50mM Tris-HCl buffer with 1.0 mM EDTA, pH 8.2 and 2.0 mM pyrogallal (in 50mM HCl). The rate of autoxidation was measured from the increase in absorbance at 420 nm and the values were expressed as U/ml.

### Catalase activity

The activity of catalase in serum of HF and HC samples was measured by H_2_O_2_ decomposition kinetics using a spectrophotometer [31]. A mixture containing potassium phosphate buffer (50mM, pH 7.0), 20mM H_2_O_2_ and 2.0μl serum was used to measure the rate of decrease in optical density at 240nm. Catalase activity was expressed as U/L.

### Glutathione reductase activity

Glutathione reductase activity was assayed by monitoring NADPH oxidation linked to GSSG reduction [32] using a spectrophotometer. The assay was performed by suspending 10μl of the serum in sodium phosphate buffer (0.3M, pH 6.8), 25mM EDTA, 20mM GSSG, 2mM NADPH and measuring the decrease in absorbance at 340 nm over time. The oxidation of 1 μmol of NADPH/min was used as a unit of glutathione reductase activity. Glutathione reductase activity was expressed as U/L.

### Glutathione peroxidase activity

Glutathione peroxidase (GPx) activity was measured by a spectrophotometric assay using 5, 5’-dithio-bis (2-nitrobenzoic acid) (DTNB) according to [33] where the oxidation of GSH in the presence of H_2_O_2_ takes place, and the unused GSH reacts with DTNB whose absorbance at 412 nm was measured. The enzyme assay was performed by incubating the serum (25 μl) in sodium phosphate buffer containing sodium azide (10mM), GSH (4mM) and 2.5 mMH_2_O_2_ for 5 minutes. Protein was precipitated using 10% trichloroacetic acid, centrifuged at 5,000 rpm for 5 minutes, and the supernatant was assayed. This assay was carried out in a microtitre plate and the readings were measured using multimode reader (Varioskan Flash, Thermoscientific, USA) after mixing with disodium hydrogen phosphate (0.3M, pH 7.0) and 100mM DTNB (in 1% sodium citrate) at 412nm. GPx activity (U/L) was measured as the amount of GSH consumed/min/mg protein after reaction with GPx and DTNB.

### Protein estimation

Serum proteins were estimated using Bradford method. A known volume (5 ul) of 1:100 diluted serum was incubated in Bradford reagent (#18004246723, Biorad, USA) and measured the absorbance at 595nm. The absorbance was extrapolated with appropriate standard reference curve using bovine serum albumin (0.031- 0.5 mg/ml) and the protein concentration was calculated. Specific activity for the antioxidant enzymes were measured after normalizing with the respective protein concentration.

### Statistical analysis

For comparing two groups, Mann – Whitney U-test was used for non-normal data and For groups with normal distribution, group means ± standard deviation (SD) were compared, homogeneity variance test (Bartlett’s test) followed by one way ANOVA was carried out. A p> 0.05 was considered to be statistically significant. For data which was not normally distributed, a nonparametric test, Kruskal-Wallis test was carried out to obtain the p value. All the statistical analyses were carried out using GraphPad Prism software version 5.0 Windows (GraphPad Software, USA).

## Results

### Stratification of heart failure patients based on circulatory redox status

Circulatory redox state (CRS) was assessed by quantifying the levels of reduced glutathione (GSH), a ubiquitous, small molecular antioxidant redox couple and malondialdehyde (MDA), an end-product of lipid peroxidation (Fig 1). Compared to the healthy controls (HC; n=42), circulating GSH levels were unchanged in HF patients (n=54) (0.09207±0.1215 μM in HF patients vs. 0.06374±0.04887 μM in HC; p=NS; Fig 1A), suggesting a wide range of redox states among the HF patients. In addition to total glutathione levels, the ratio of reduced to oxidized glutathione (GSH:GSSG) did not differ between HF and HC (4.93±7.25 in HF patients vs. 3.01±1.78 in HC; p=NS; Fig 1C). In contrast, MDA was significantly higher in HF patients vs. HC (4.342±2.103 vs.1.566±0.3018 μM, respectively; p<0.0001) (Fig 1B), suggesting that increased lipid peroxidation is coupled with the development of HF. Therefore, we next assessed whether the redox index (reduction vs. oxidation) is significantly different between the groups. Comparisons of GSH/GSSG ratio revealed insignificant differences between the groups, indicating that the HF patients seem to exhibit a wide range of redox levels in relation to HC (4.93±7.25 in HF patients vs. 3.01±1.78 in HC; p=NS; Fig.1C). Therefore, our approach for stratifying HF patients used CRS (i.e. GSH/MDA) to gain more knowledge and understand the pathogenesis. But, comparisons of GSH/MDA ratios between the HF and HC yielded a statistically significant result (0.02±0.0345 in HF patients vs. 0.04±0.028 in HC; p<0.0001; Fig.1D). Therefore, we were interested in stratifying the HF patients based on their circulating redox score (CRS) (Fig 1E). The CRS of healthy subjects was designated as normal redox (NR). A CRS value in any patient lower than the NR displayed by HC was regarded as hyper-oxidative (HO) while a patient exhibiting a GSH/MDA ratio greater than the NR cutoff was classified as hyper-reductive (HR). Surprisingly, our CRS-based stratification revealed 3 distinct classes within our cohort of HF patients (Fig. 1E). Along these lines, 41% of HF patients were assigned to the normal redox (NR) group whose mean GSH/MDA ratio (0.02335±0.01201) was similar to HC (0.03969±0.02857). Next, 42% of HF patients belonged to the hyper-oxidative (HO) class whose mean GSH/MDA ratio (0.004787±0.002055) was lower than HC group (0.03969±0.02857), and 17% of HF patients exhibited a hyper-reductive (HR) state in which mean GSH/MDA ratio (0.2271±0.03192) was greater than HC group (0.03969±0.02857). Interestingly, the GSH/MDA ratios were statistically significant among all three HF groups (p<0.0001) by one way ANOVA analysis. These novel observations suggest that redox status plays a variable role in the progression of HF and that both extremes (e.g. oxidative and reductive) may contribute to disease pathogenesis.

**Figure 1.**
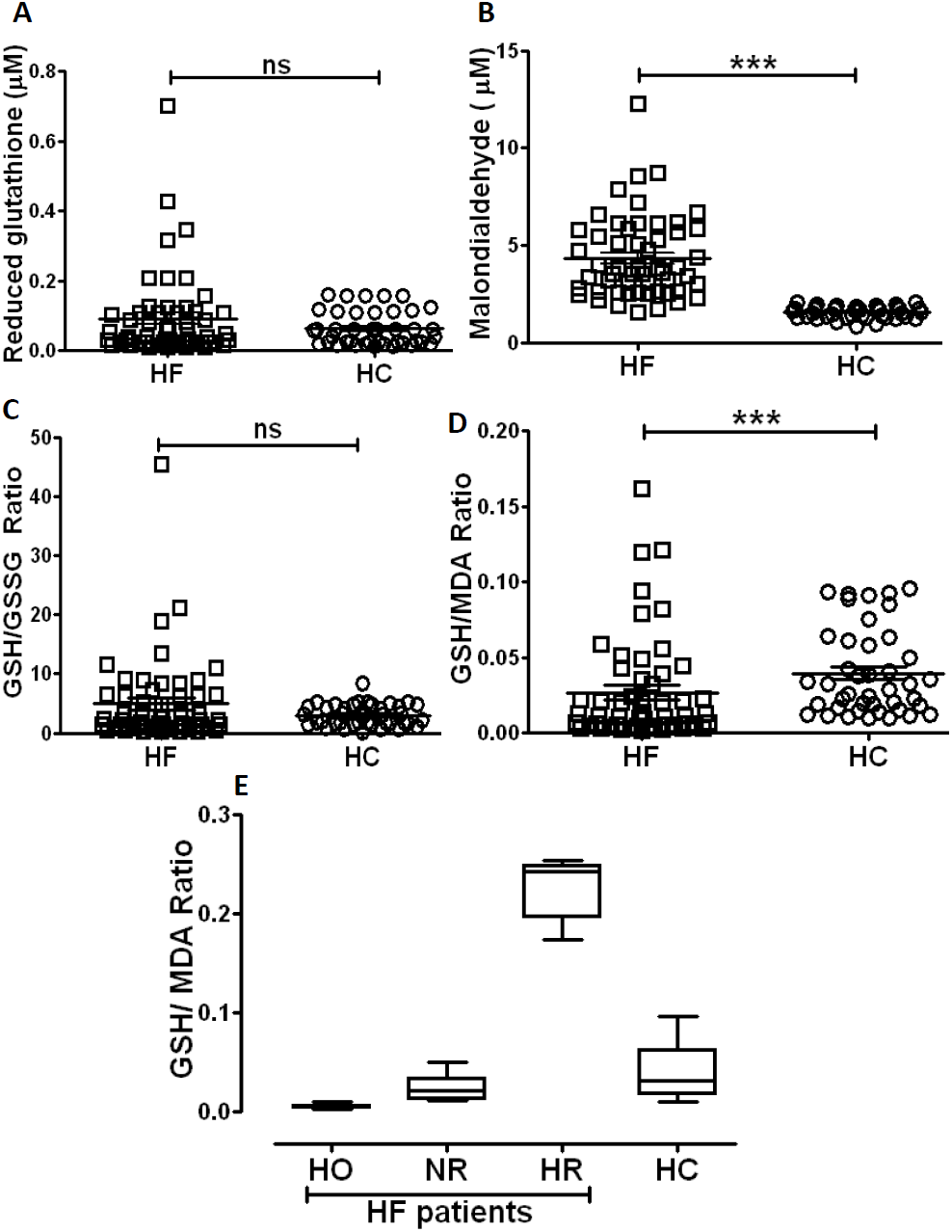
Stratification of heart failure (HF) patients based on redox status. A. Comparison of reduced glutathione (GSH) levels showing statistically insignificant GSH levels between HF patients and HC. B. Significantly higher malondialdehyde levels in HF patients vs. HC. C. Statistically insignificant GSH/GSSG ratio between HF patients and the healthy controls. D. Mann – Whitney U-test was carried out for statistical significance. ^***^p < 0.001. n=42 HC and 54 HF patients. E. Stratification of HF patients into hyper-oxidative (HO), normal redox (NR) and hyper-reductive (HR) groups based on comparison of GSH/MDA ratio with healthy controls (HC). Remarkably, higher GSH/MDA ratio in HR group, similar GSH/MDA ratio in NR and HC, and a significantly lower GSH/MDA ratio in HO group were observed. One way ANOVA was carried out with p<0.0001 among all the heart failure patient groups (n=42 HC, 23 HO, 22 NR, and 9 HR).

### Activities of antioxidant enzymes were significantly altered in HF vs. healthy controls

To assess the capacity of the endogenous antioxidant defense system in the periphery of HF patients, we further analyzed the activity of circulating antioxidant enzymes such as superoxide dismutase (SOD), glutathione reductase (GR), catalase, and glutathione peroxidase (GPx) (Fig. 2). While the activities of SOD (6.566±2.297 vs. 2.571±0.6179 U/ml; p<0.0001; Fig. 2A), GPx (490.4±299.6 vs. 312.0±141.4U/L; p<0.001; Fig 2B) and catalase (864.0±622.2 vs. 689.1±556.8 U/L; p<0.05; Fig. 2C) were significantly increased, GR activity was significantly lower (13±7.648 vs. 24.70±28.22 U/L; p<0.05; Fig. 2D) in HF patients relative to healthy controls.

**Figure 2.**
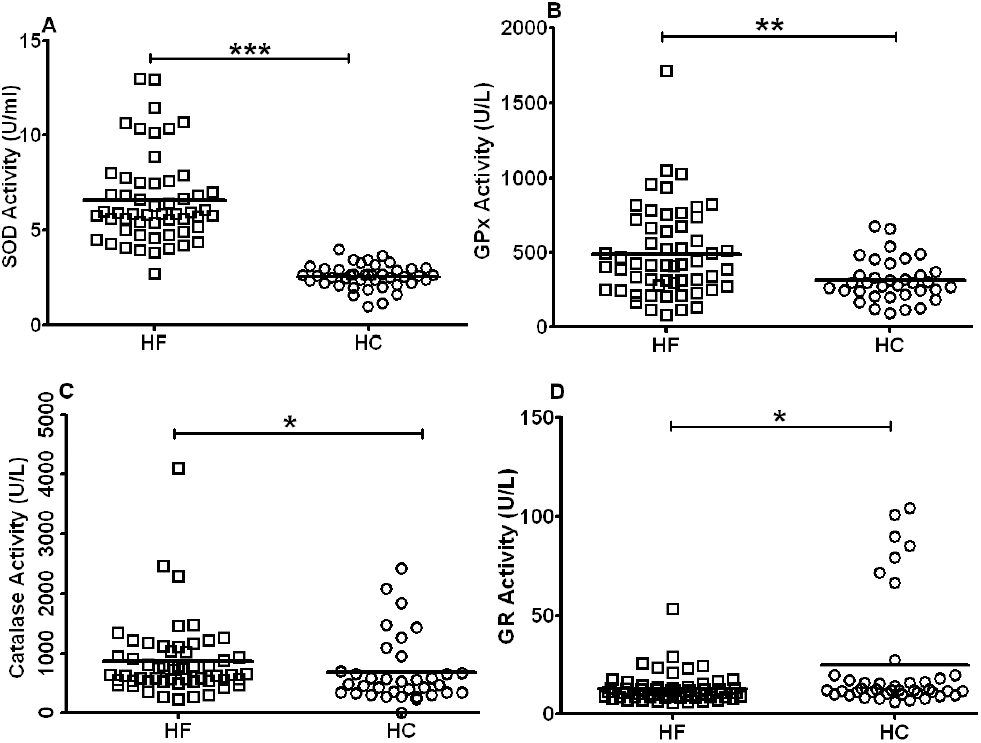
Comparison of antioxidant enzymes between heart failure (HF) patients and healthy controls. A. Statistically significant higher superoxide dismutase activity(SOD) in HF patients compared to healthy controls. B. Statistically significant higher glutathione peroxidase activity (GPx) in HF patients compared to healthy controls. C. Statistically significant higher catalase activity compared to HC. D. Mann – Whitney U-test was carried out for statistical significance. ^***^p<0.0001, ^**^p<0.01, ^*^p<0.05. n = 42 HC and 54 HF patients, except for catalase and GPx, where n=35 HC because of limited sample volume.

### Redox associated changes in cardiac remodeling and systolic function in HF patients

To relate our biochemical redox observations in the distinct subsets of HF patient with myocardial structural and functional remodeling, we performed non-invasive echocardiography analyses to determine fractional shortening, ejection fraction and left ventricular (LV) mass. First, we confirmed that HF patients exhibit significantly distinct myocardial mass and systolic function when compared to HC (Fig. 3A-C). Fractional shortening (Fig. 3A), ejection fraction (Fig. 3B) and LV mass (Fig. 3C) did not show any statistically significant differences among NR, HO, and HR groups of HF patients. However, when compared with HC, all HF patient groups (HO, NR and HR) showed significantly lower values for fractional shortening (17.03±9.256 for HO, 15.40±5.863 for NR, 16.54±5.185 for HR vs. 36.13±6.020 for HC; p<0.01) and ejection fraction (33.78±17.62 for HO, 32.47±10.41 for NR, 32.57±9.424 for HR vs. 65.46±7.9 for HC; p<0.01) but significantly greater LV mass (309.1±107.9 for HO, 281.1±128.2 for NR, 320.1±103.8 for HR vs. 162.6±13.72 for HC; p<0.01). Representative M-mode echo images for systolic function analysis of NR, HO, HR and healthy control groups are shown in Fig 3D, 3E, 3F and 3G, respectively.

**Figure 3.**
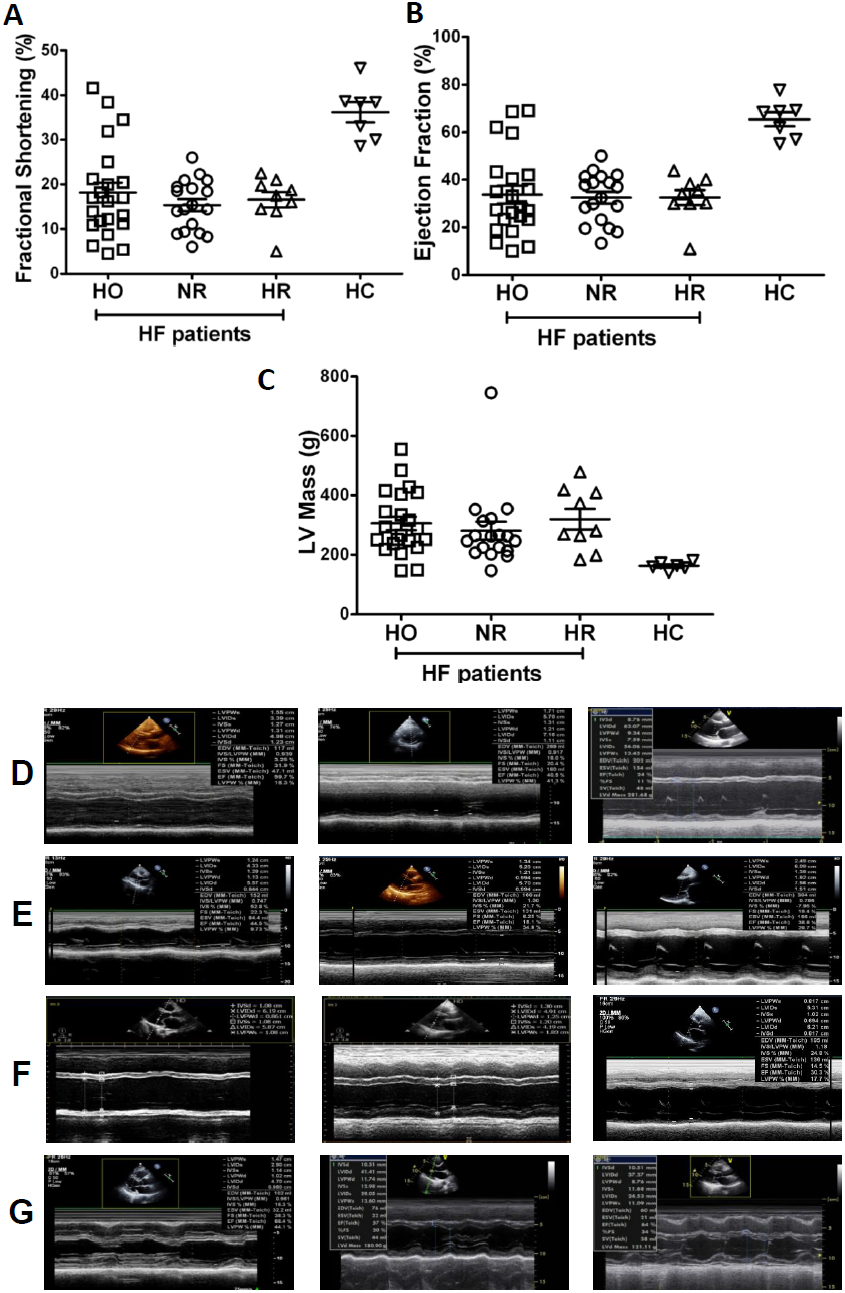
Echocardiographic parameters for hyper-oxidative (HO), normal redox (NR), hyper-reductive (HR) and healthy controls (HC). A. Statistically significant lower percentage of fractional shortening (FS) for HO, NR and HR groups of HF patients compared to HC. B. Statistically significant lower percentage of ejection fraction for HO, NR and HR groups of heart failure patients compared to HC and C. Statistically significant higher LV mass for HO, NR and HR groups of heart failure patients compared to HC. One way ANOVA was carried out with a p<0.01 vs. HC for A, B and C. n = 7 HC, 22 HO, 18 NR and 9 HR. M-mode ECHO related parameters focusing on systolic functions and dimensions with ejection fraction (EF) and FS for D. HO, E. NR, F. HR and G. HC. n=3 representatives/group were presented.

### Evidence for a wide range of diastolic abnormalities in HF patients

Primary measurements of trans-mitral inflow including the peak early filling (E wave), late diastolic filling (A wave) velocities and the E/A ratio were performed in HF patients as well as HC (Fig. 4A-C). Although there was no statistically significant difference between HF patients and the HC for ME (88.91±29.67 for HO, 91.19±37.26 for NR, 81.26±33.67 for HR vs. 75.64±18.60 for HC; p=NS), MA (71.47±30.53 for HO, 59.65±25.25 for NR, 58.44±21.22 for HR vs. 65.41±11.65 for HC; p=NS) and ME/A ratio (1.50±0.92 for HO, 1.76±0.97 for NR, 1.66±1.12 for HR vs. 1.18±0.34 for HC; p=NS), their values were widely distributed among the groups compared to the controls (Fig. 4D-F). Representative mitral valve Doppler analysis for diastolic function with graphs for ME/A ratios for HO, NR, HR and healthy control groups are shown in Fig 4G, 4H, 4I and 4J respectively. M E/A ratio differed according to age and gender [34, 35]. Based on this range, we grouped the HF patients into impaired relaxation and restrictive filling.

**Figure 4.**
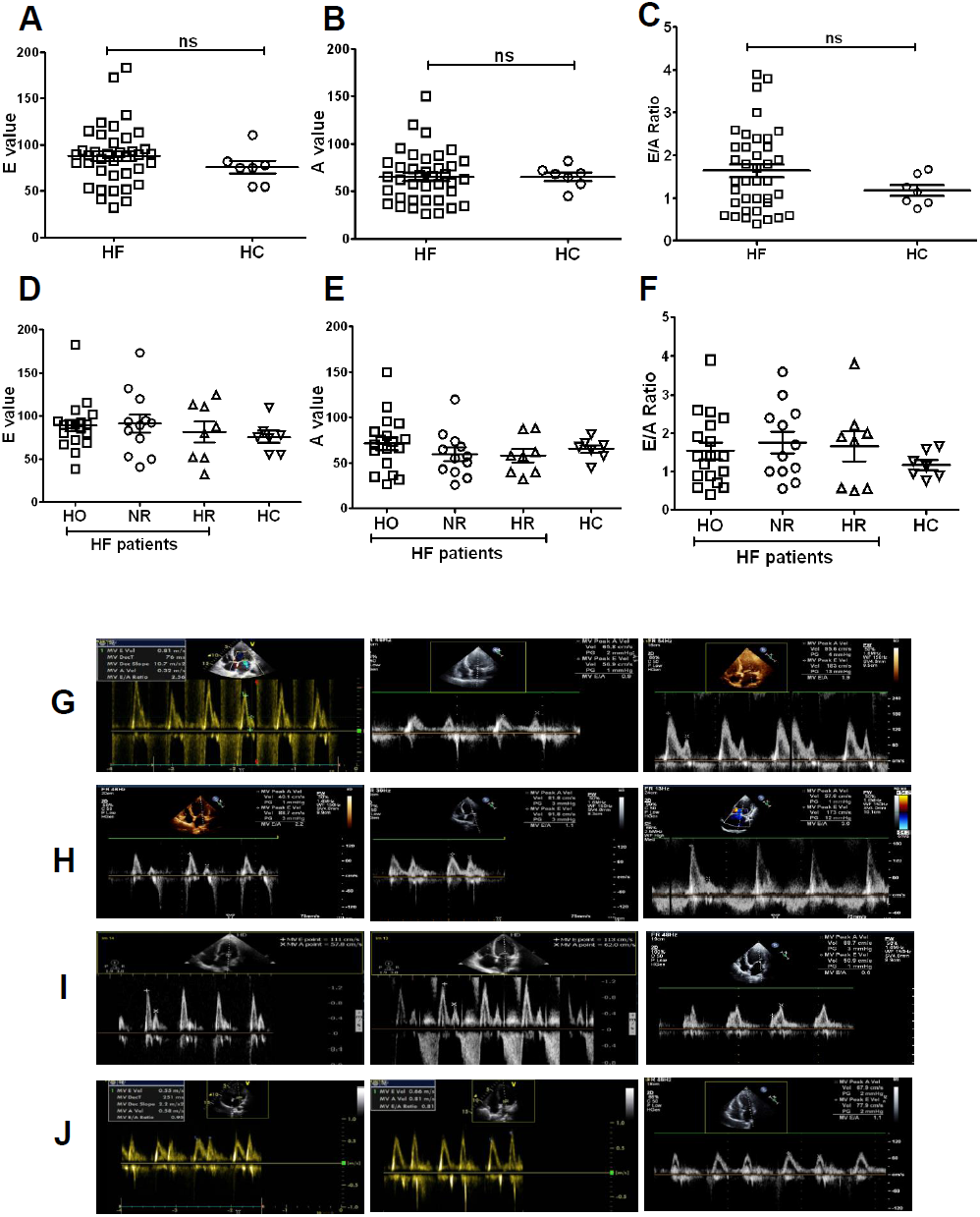
Comparison of diastolic function abnormalities between HF patients and healthy controls. Similar values between HF patients and healthy controls were observed for A. Early (E) filling, B. late (A) filling and C. E/A ratio. Mann – Whitney U-test was carried out with no statistical significance among the groups for A, B and C, n = 7 HC and 38 HF patients. D. Early (E) filling values were slightly higher in HF group patients (HO, NR and HR) than healthy controls. Relative to healthy controls, E. late (A) filling values and were slightly higher in HO and lower in NR and HR. F. E/A ratio were slightly higher in HF group patients (HO, NR and HR) than healthy controls. One way ANOVA was carried out with no statistical significance among the groups for D, E and F. Mitral valve Doppler analysis for diastolic function with graphs for M E/A ratios for G. Hyper-oxidative, H. Normal redox, I. Hyper-reductive and J. Healthy controls.

### Proportion of patients based on systolic (EF) and diastolic (E/A ratio) functions suggest a hyper-reductive condition may be detrimental factor in HF pathogenesis

Among the HF patients, the HO group (55%) had a higher percentage of patients with ejection fraction ≤ 30% than the NR (44.5%) and HR groups (44.5%). Patients with an EF between 31 – 40% were more frequently assigned to the HF group (33.3%) compared to NR (28%) and HO groups (18%). In contrast, HF patients displaying an EF between 41 – 50% were less prone to be classified as HO (9%) when compared to the NR (22%) and HR groups (22%). Intriguingly, HF patients with an EF≥50% were completely absent in HR groups relative to HO (18%) and NR patients (5.5%) (Fig. 5A).

**Figure 5.**
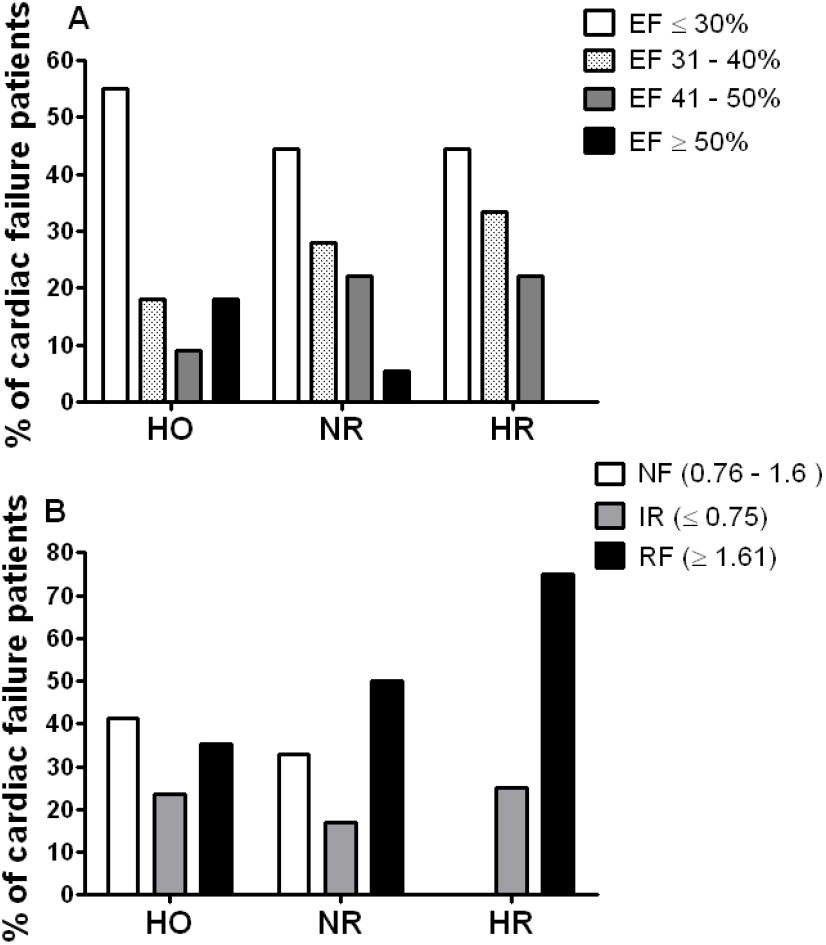
Comparison of systolic functions (EF) and diastolic functions (E/A ratio) among hyper-oxidative, normal redox and hyper-reductive groups of HF patients. A. Comparison of ejection fraction (EF) among hyper-oxidative (HO), normal redox (NR) and hyper-reductive (HR) groups of HF patients indicating higher number of HO patients have lowest range of EF ≤ 30% and patients with highest range of EF ≥ 50% are absent in HR group. B. Comparison of E/A ratio among HO, NR and HR groups of cardiac failure patients with normal filling (NF), impaired relaxation (IR) and restrictive filling (RF) indicating that the HR group had higher number of patients with RF and HO group had higher patients with NF.

Using the ME/A ratio as an index, diastolic function was compared between different HF groups and healthy controls (Fig. 5B). To our surprise, none of the HR patients exhibited normal filling (E/A ratio 0.76 – 1.6); however, 41.3% of HO and 33.3% of NR patients revealed such E/A ratios. While only 17% of NR patients exhibited impaired relaxation (E/A ratio ≤ 0.75), the incidence of impaired relaxation in HR (25%) and HO (23.5%) patients. was evenly distributed. Finally, the HR group had higher percentage of patients (75%) with restrictive filling (E/A ratio ≥ 1.61) as compared with HO (35.2%) and NR patients (50%).

## Discussion

The current understanding of redox changes during heart failure (HF) is centered on results obtained for a given group of HF patients compared to healthy controls. Importantly, this experimental approach assumes that redox imbalances during HF progression confirm to the oxidative stress paradigm and neglect the potential for observations of individual redox responses in unique subsets of cardiac patients. Here, we attempted to investigate the individual differences of circulating redox biomarkers such as reduced glutathione (GSH), its redox ratio (GSH/GSSG), lipid peroxidation levels (i.e. MDA) and its normalization to reduced glutathione (i.e. GSH/MDA). In particular, we examined whether some HF patients exhibit a unique redox state that is likely to be stretched on either direction of the redox spectrum (i.e. hyper-reductive vs. hyper-oxidative). The development of HF might occur in response to various factors including a chronic oxidative stress/inflammation, infection, ischemic insult, genetic cues and other chronic stresses [36, 37]. Hither to, the redox-based classification of HF has not been implemented.

Generation of reactive oxygen/nitrogen species (ROS/RNS) and oxidative stress has been reported to be associated with the development of HF [38, 39]. However, the evidence for oxidative stress as a causal factor for chronic HF is ambiguous as oxidative stress is absent at the onset of HF in some patients while it appears as a consequence at the later stages of the disease in others. Therefore, we postulate that the oxidative stress may not be solely responsible for the development of HF, and that the other extreme of the redox continuum, reductive stress (RS; a hyper-reductive condition) may play an equally pathological role in certain instances. In the past decade, we and others have reported that RS might also be an additional mechanism that causes pathological cardiac remodeling leading to heart failure [28, 40-42]. Notably, the existence of RS as a key driver of pathogenesis has been shown in a human mutant protein aggregation cardiomyopathy with augmented antioxidant capacity [28, 43]. Furthermore, other laboratories have reported that augmentation of HSP25, a molecular chaperone that co-regulates glutathione metabolism, promoted RS and resulted in cardiac hypertrophy in a transgenic mouse expressing Hsp25 [42]. Of late, it has been shown that significant upregulation of NADPH in the heart promotes reductive stress and exacerbates myocardial injury in a mouse model [44]. Nonetheless, such an evidence for RS or a hyper-reductive condition in human HF has not been demonstrated. Our current observations in a small cohort of HF patients (n=54) have indicated that a distinct subset of HF patients exhibit a hyper-reductive condition in their circulation. Thus, we believe that our novel findings might be useful in stratifying HF patients based on the circulatory redox state (CRS) and design appropriate treatment strategies targeting antioxidants.

Based on the CRS (i.e. GSH/MDA), HF patients were stratified into 3 distinct groups, namely HF patients with (a) normal redox (NR), (b) hyper-oxidative (HO) and (c) hyper-reductive (HR) conditions. Interestingly, while a majority of the HF patients fall into the HO (42%) and NR (41%) categories, a small subset exhibited the HR (n=17%). Of note, these observations suggest that not all HF patients experience oxidative stress. Therefore, a key question of how and why NR or HR redox conditions contribute to the development of HF remains. Although these analyses seem to be novel and interesting, the mechanisms for NR and/or HR redox conditions in the development of HF are yet to be discovered. Importantly, changes in circulating redox biomarkers in a single group of HF patients appear to be insignificant when compared to healthy controls as the individual levels of CRS have a wide range, which masks the actual scenario in a given HF patient. Hence, our approach for stratifying HF patients based on their CRS will be appropriate to gain more knowledge and understand the pathogenesis of HF in a personalized manner that is likely to enhance the potential of a patient’s treatment. Furthermore, we believe that a basic and simple biochemical clinical test for the CRS of a HF patient is a feasible strategy to improve the patient’s health and increase their survival through selecting appropriate treatment strategies.

In the majority of the patients (>85%), our CRS measurements were performed upon the first hospital admission. Critically, we meticulously confirmed that no subjects had consumed antioxidant supplements prior to this visit, thereby indicating a clear picture of a hyper-reductive condition associated with HF. In addition to providing the first evidence for an abnormally reductive state in human HF patients, another key finding of our study is the wide heterogeneity of the CRS in response to HF. Along these lines, the prevalence of NR and HR in a considerable number of participants contradicts the expected HO. Considering that ROS/RNS serve as key cellular signaling molecules for basal physiology and responses/adaptations to acute/chronic stress conditions [45-47], patients who exhibit NR or HR may have not gained protection from developing HF. Altogether, these results indicate that in contrast to the common understanding, oxidative stress is not the only factor inducing HF.

Next, our detailed analyses of cardiac structure and function among the 3 classes of HF patients revealed interesting information. Most of the HO patients displayed severe systolic dysfunction with an ejection fraction ≤ 30% while comparing with NR and HR group. It has been established that an over production of ROS/RNS adversely alters cardiac function eventually leading to diastolic and systolic dysfunction [48, 49]. Moreover, left ventricular (LV) dysfunction in HF patients correlates well with the extent of oxidative stress in the myocardium and plasma [48]. In HR group none of the patients have ejection fraction ≥ 50% and only 5% in NR group. Hence both HR and NR groups predominantly have mild to moderate systolic dysfunction. As such, we speculate that oxidative stress could be a major cause for severe systolic dysfunction rather than reductive stress. In addition to systolic function, we assessed the grade of diastolic dysfunction among the three groups of HF patients utilizing noninvasive Doppler echocardiography mitral inflow (E/A ratio). Surprisingly, the HR group exhibited a higher percentage of patients with a restrictive filling pattern while nil for normal filling pattern. However, all other groups had almost equal percentage of patients with normal filling pattern. From our data, it is apparent that patients with RS have severe diastolic dysfunction as compared with normal redox and oxidative stress conditions. Specifically, reductive stress imparts both mild to moderate systolic dysfunction and severe diastolic dysfunction among HF patient groups. Up-to-date, there has been accumulating evidence indicating that HF with systolic dysfunction is associated with oxidative stress and nitric oxide (NO) signaling [50] whereas our observations suggest a novel role for reductive stress in the development of diastolic dysfunction.

## Conclusion

In this study, we believe that we provided evidence supporting the presence of a hyper-reductive (i.e. reductive stress) condition in a subset of HF patients. Notably, we observed that HR HF patients had diastolic dysfunction while HO HF patients exhibited systolic dysfunction. Further, individual differences in CRS during the development of HF are apparent suggesting unique contributions of redox disequilibrium across distinct classes of patients. The exact nature of the mechanisms responsible for the heterogeneity in redox responses in the context of HF is presently unknown. We consider that the wide inter-individual variability for CRS shown here is not limited to the two biomarkers (GSH and MDA) of the redox milieu. Remarkably, the data presented herein emphasize that the mean CRS of a group of HF patients can be misleading. We also acknowledge that this pilot study included a small (but tightly controlled for age and other comorbidities) and male dominant cohort of HF patients. Therefore, our future investigations will focus on larger groups including both genders with specific types of HF such as ischemic disease, hypertrophic cardiomyopathy, dilated cardiomyopathy, and rheumatoid cardiac diseases, and will emphasize differences in diastolic vs. systolic dysfunctions.

## Acknowledgements

This study was supported by funding from PSG Center for Molecular Medicine and Therapeutics, PSG Institute of Medical Sciences & Research (For SR), Coimbatore, Tamil Nadu, India and Department of Pathology/School of Medicine startup funds (for NSR Travel) University of Alabama at Birmingham, AL.

The authors greatly appreciate Drs. Anil Challa and Shanmugam Gobinath, and Mr. Justin Quiles for preview and editorial assistance.

